# Simulation atomic force microscopy to predict correlated conformational dynamics in proteins from topographic imaging

**DOI:** 10.1101/2021.10.15.464530

**Authors:** Holger Flechsig

## Abstract

Atomic force microscopy (AFM) of proteins can detect only changes within the scanned molecular surface, missing all motions in other regions and thus information about functionally relevant conformational couplings. We show that simulation AFM can overcome this drawback by reconstruction of 3D molecular structures from topographic AFM images. A proof of principle demonstration is provided for an in-silico AFM experiment visualizing the conformational dynamics of a membrane transporter. The application shows that the alternating access mechanism underlying its operation can be retrieved from only AFM imaging of one membrane side. Simulation AFM is implemented in the freely available BioAFMviewer software platform, providing the convenient applicability to better understand experimental AFM observations.

## Introduction

Atomic force microscopy (AFM) and high-speed AFM scanning (hsAFM) allow to visualize structures of biomolecules and monitor their functional conformational dynamics under near-physiological conditions (see the reviews^1,2^). A major limitation is that AFM scanning detects only morphological changes within the probed molecular surface, missing all motions in other regions and thus information about functionally relevant couplings of conformational dynamics.

A representative example is that of a transporter protein (Fig. 1A) which performs cyclic conformational changes, driven e.g. by the energy from ligands, to mediate the transmembrane exchange of substances. Their operation often involves switching between an inward-facing conformation (open to the intracellular side) and an outward-facing conformation (open to the extracellular), referred to as the alternating access mechanism^3^ (Fig. 1B). Scanning either side of the protein does not allow to explore intramolecular communication underlying transport and explain such a mechanism (Fig. 1A). Even if two independent experiments would be conducted, scanning the protein surface from both sides, it would still remain impossible to capture the synchronized behavior.

**Figure 1.**
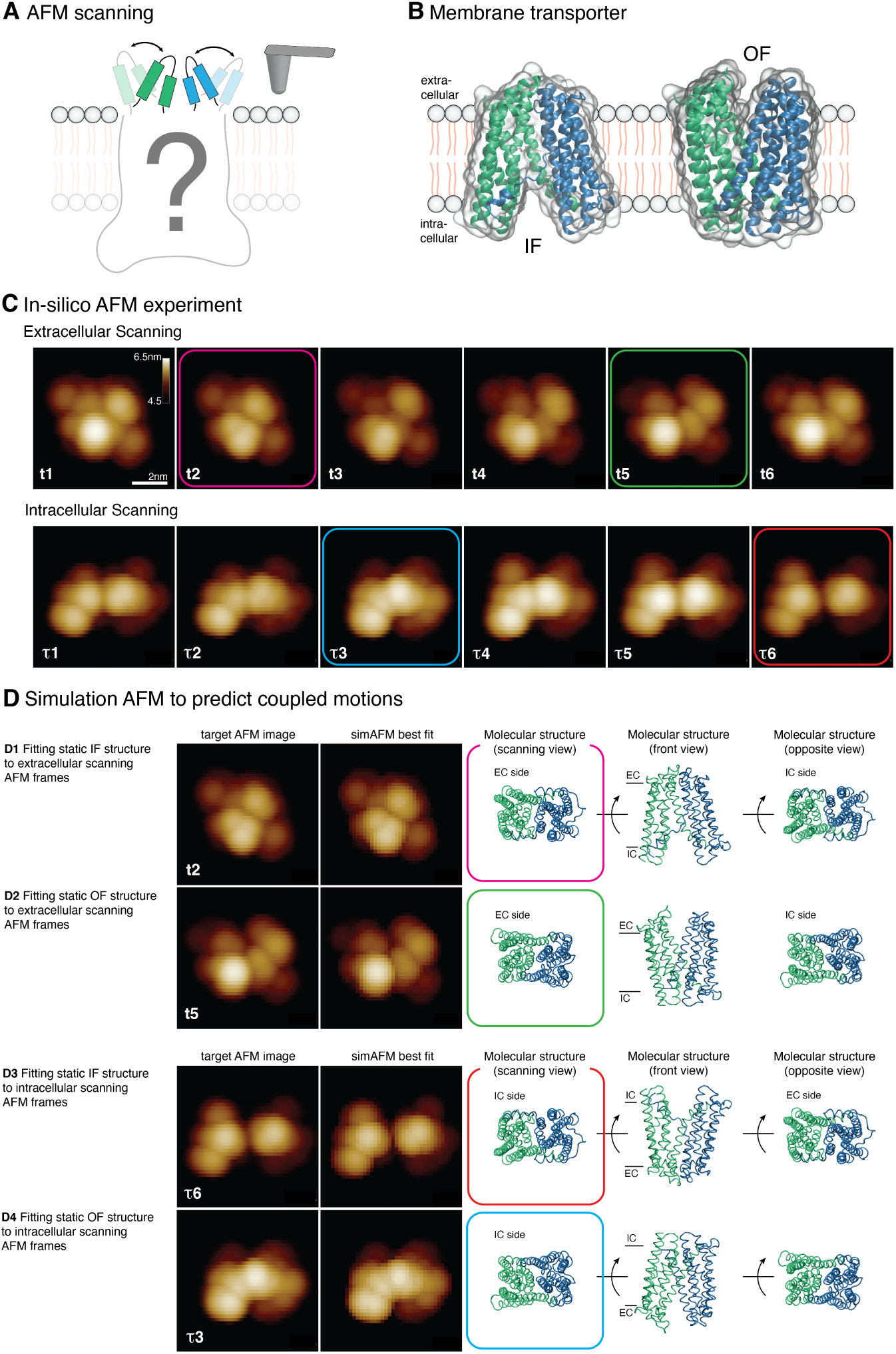
A: AFM scanning of a membrane protein, detecting motions only at one side. B: Static structures of the PfMATE transporter in the inward-facing (IF, 6FHZ) and outward-facing (OF, 4MLB) conformation. C: Selected topographic images from the in-silico PfMATE AFM experiment at different time moments for scanning from extra- and intracellular sides. Images are filtered by a motion blur in the scanning direction. D: Simulation AFM of PfMATE IF and OF static structures and fitting to AFM movie frames. Best matches of IF (D1) and OF (D2) structure to frames of extracellular scanning. Corresponding molecular structures are shown in scanning and side view, and from the opposite side. D3,D4: Best matched of IF/OF structures to frames of intracellular scanning and corresponding retrieved transporter structures. Colored frames mark the pairs of AFM images with their corresponding reconstructed molecular structures.

Generally, the drawback is present in AFM experiments of protein machines and motors which undergo correlated conformational dynamics coupling the motions of remote regions during their functional activity.

While this intrinsic limitation of AFM can in principle be overcome to some extend, e.g. by a combination with optical or fluorescent spectroscopy methods (e.g.^4^), the experimental realization is highly challenging. On the other side, computational approaches including molecular modelling and simulations are inreasingly used to assist in understanding biomolecular high-speed AFM measurements^5–8^.

We recently developed the BioAFMviewer software platform, which implements simulation atomic force microscopy (simAFM) of biomolecular structures and molecular movies of their conformational dynamics^9^. We further demonstrated how simAFM and automatized fitting to experimental images allow to reconstruct 3D atomic structures from AFM surface scans and enable quantitative molecular feature assignment within measured topographies ^10^.

In this short report we show that simulation AFM can be employed to predict conformational couplings in proteins from topographic AFM images, thus opening a novel perspective on its applications to facilitate the interpretation of AFM measurements.

## Results and discussion

### In-silico AFM experiment

We performed an in-silico AFM experiment of the *multidrug and toxic compound extrusion* (MATE) transporter from *Pyrococcus furiosus* (PfMATE). For this transporter the atomic structures of inward-facing (IF) and outward-facing (OF) conformations are known (see Figure 1B). Using them as references states, the conformational dynamics underlying the transition IF *→* OF *→* IF was computed in elastic-network based coarse-grained Brownian dynamics simulations implemented in the eBDIMS software^11^. Thus, a molecular movie of feasible conformational motions along the operation cycle of the MATE transporter was obtained (Supplementary Movie 1). This movie was processed by the BioAFMviewer software which performs simulation AFM to produce a computerized AFM experiment^9,10^. We conducted two in-silico AFM experiments of the MATE transporter operation cycle, scanning the extracellular and the intracelluar side, respectively. Corresponding AFM movies are provided as Supplementary Movie 1. A series of simAFM topographic images is shown in Figure 1C. The complete set of simAFM images from each in-silico AFM experiment of PfMATE was used as a template for our demonstration.

### Simulation AFM to predict intramolecular communication

For a proof of principle demonstration the following situation is considered. We have AFM movies from two independent in-silico experiments. For each topographic AFM image the molecular conformation of the transporter underlying the obtained scan and information about its instantaneous structure at the respective other side of the membrane is not known. Then, the following strategy was applied. The atomic PDB structures of PfMATE in the static IF (6FHZ) and OF (4MLB) conformation served as templates for which simulation AFM was performed within the BioAFMviewer software interactive interface. For either template structure, the automatized fitting function identified the best match of the simAFM image with a target AFM image of PfMATE. The target images consisted of the complete set of frames from both AFM movies.

First we focused on the AFM experiment from the extracellular side. Performing fitting of the IF structure individually to all 200 AFM movie images gave the best match of simAFM and target AFM image for the snapshot at time moment *t*2 (Fig. 1C and D1), with an image RMSD of 1.73. Therefore, the full information about the molecular structure of the transporter for this AFM topographic image is obtained (shown in scanning, side, and opposite view orientations in Fig. 1,D1). The AFM image corresponds to a transporter conformation which is closed to the extracellular (scanning) side, while its overall V-shape causes the intracellular side to be wide open. For fitting of the OF structure the overall best match was obtained with the AFM movie frame at time moment *t*5 (Fig. 1C and D2). SimAFM and target AFM image compared with an RMSD of 4.98. The retrieved molecular structure, shown in different orientations in Fig. 1D2, reveals that this particular AFM image monitors the transporter in a conformation wide open to the extracellular (scanning) side, whereas its instantaneous conformation at the opposite membrane site is shut to the intracelullar side.

Hence, it is demonstrated that from only an AFM image of the transporter extracellular surface, the full 3D information of its functional conformation, including intramolecular coupling of domain motions, can be obtained.

We repeated the procedure for the AFM experiment of the intracellular side (Fig. 1 D3,D4). The IF transporter structure matched best to the AFM frame at time *τ*6 (RMSD 3.85), identifying its topography to represent the molecular structure with an opened conformation whose corresponding opposite extracelluar side is closed (Fig. 1 D3). The best match of the OF structure was obtained for the AFM image at *τ*3 (RMSD 4.41), which therefore represents the open conformation and predicts the opposite side to have the closed state.

The application of simulation AFM to the in-silico AFM experiment of a membrane transporter shows that the alternating access mechanism underlying its operation can be retrieved from AFM imaging of only one membrane side.

## Discussion

We show that simulation AFM can be employed to predict correlated conformational dynamics in proteins from topographic AFM images. The application was demonstrated for an in-silico AFM experiment of a transporter protein, showing that the alternating access mechanism underlying its operation can be retrieved from imaging only the topographical changes at one membrane side.

Although transmembrane channels and transporters are widely investigated with high-speed AFM^12^, a study which could visualize significant motions at both membrane sides is not available (to our knowledge). Hs-AFM imaging of bacteriorhodopsin showed conformational motions at the cytoplasmic side upon photo-activation, but not at the extracelluar side^13^. Similarly, a study of an ion channel evidenced nucleotide-dedendent motions at the intracellular side, while a clear coupling to changes at the extracellular side could not be obtained^14^. It should be expected that AFM imaging of active transporters, whose operation typically involves large-scale domain motions coupling opposing structural regions, leads to exciting new insights about the transport mechanism. Multidrug efflux transporters or ATP-binding cassette (ABC) transporters can be targets of interest. If such experiments are realized, simulation AFM provides the framework to better understand experimental observations.

Simulation AFM is implemented in the freely available BioAFMviewer software, providing the convenient platform for applications to experimental AFM data.

## Acknowledgement

This work was supported by the Ministry of Education, Culture, Sports, Science and Technology (MEXT), Japan, through the World Premier International Research Center (WPI) Initiative, and by Japanese Society for Promotion of Science Grant-in-Aid for Scientific Research 21K03483.

